# Genetic Variation in *Drosophila melanogaster* Aggression

**DOI:** 10.64898/2026.07.04.736468

**Authors:** Jennifer M. Gleason, Cailin M. Kessen, Vaishnavi Verma, Eleanor Bath

## Abstract

Animals fight for resources to obtain fitness benefits; most contests are intrasexual, and males tend to fight more than females. Although the genetic basis of male aggression is well studied, we know little about the genetic variation of female aggression. Female aggression varies with reproductive status and is potentially influenced not only by her genotype, but also by the genotype of her mate. Here we measured both male and female aggression in a set of *Drosophila melanogaster* inbred lines by competing each line against a standard competitor. Aggression varied among lines for both sexes, but male and female aggression were not correlated. Female aggression for many lines increased with mating, as expected, but not all lines changed aggression. However, when females were mated to males of different lines, male genotype did not affect the post-mating change in aggression, suggesting that ejaculate-mediated effects do not vary across these lines. The aggression level of the standard opponent was positively correlated with that of focal individuals indicating that individuals modulate their behavior according to the genotype of their opponent.

## Introduction

### Aggression

Contests among conspecifics of the same sex may determine an individual’s access to resources, including territory, food, and mates (1). Winning aggressive encounters can therefore provide fitness benefits. However, aggression also comes with costs in the form of potential injury, wasted time, and energy expenditure (2). Individuals should therefore choose carefully when to engage in aggression. Despite the costs and benefits, populations vary in how much individuals fight (3). A substantial amount of variation in aggression is likely due to the environment shifting the costs and benefits of engaging in aggression. The number of competitors, the condition of potential competitors, the risk of predation, and value of the contested resource all influence rates of aggression (2).

Although aggression changes plastically in response to the environment, aggression has a genetic component. Heritability of aggression in mammals and birds is high (reviewed in 4). Aggression responds to selection: selection against aggression is common in domestication (e.g. 5, 6), though aggression readily increases in response to artificial selection as well (e.g. 5, 7, 8). The underlying genetic architecture of aggression is complex, with many genes contributing to the expression of aggression. Individual genes have been identified that contribute to aggression, though most are known from their contribution to extreme aggression phenotypes (reviewed in 4) and not for contributions to natural variation. Populations vary in levels of aggression depending on differences in selection pressures (e.g. 9-11). Population-level variation may reflect fluctuating selection on aggression as the costs and benefits of aggression are related strongly to the current social environment, or frequency-dependent selection resulting in aggressive and non-aggressive phenotypes (12).

Although genes have been identified affecting male aggression in many species, particularly humans, mice, and *Drosophila* (reviewed in 4), we know little about genetic variability of aggression in females. Females, like males, compete aggressively with fitness consequences (13). Intrasexual aggression in females has been relatively neglected, especially outside species with dominance hierarchies (14). Whereas males tend to compete over access to mates and territories, females are likely to compete over access to food, nesting sites, and resources for their offspring (13). The form, frequency, and intensity of aggression often differ significantly between the sexes (13), suggesting both differential selection and underlying differences in the genetic architecture regulating aggression. Selection for aggression may differ between males and females due to differences in contested resources and the relative costs and benefits of aggression (13). Commonalities of male and female aggression, particularly the underlying genetic basis, are relatively unexplored.

### Drosophila aggression

Both male and female *Drosophila melanogaster* fight with conspecifics. Males are the more well-studied sex, with evidence suggesting the mechanisms that regulate male aggression are highly conserved from flies to mammals (4, 15). Male *Drosophila melanogaster*, in general, are more aggressive than females (16, 17), engaging in aggression to defend territories and guard mates (18). Males have a wide repertoire of aggressive behaviors, ranging from low-intensity, non-contact behaviors (wing threats, approaching and chasing (19)) to higher intensity contact behaviors (lunging, holding, chasing, boxing, and tussling). Female-female aggression in this species is well described (e.g. 20-24) but remains understudied compared to males. Females are most likely to fight over food and oviposition sites but also attack mating pairs (25). Females have a different set of aggressive behaviors, including low-intensity behaviors (wing threats and fencing) and higher intensity behaviors (headbutts and shoving, 17).

Male aggression is heritable in *Drosophila melanogaster* (26) and differs significantly among genetic lines of *Drosophila melanogaster*, varying more than twentyfold between the least and most aggressive of 200 strains (27). Variation among strains in female aggression has been tested on a much smaller scale, with no differences found between three commonly used lab strains (21). Therefore, the range of variation in female aggression is unknown. Aggression in both sexes evolves in response to artificial selection and experimentally manipulated sex ratios, suggesting a genetic basis that can respond to selection (16, 28).

The genetic basis of aggression is not necessarily the same in males and females. Evidence for an intersex genetic correlation in aggression in *D. melanogaster* is mixed. Selection experiments on male aggression have resulted in both a correlated change in female aggression (28), and no change (29). In contrast, aggression in the two sexes had a positive correlation across populations evolved at different sex ratios, suggesting a possible shared genetic basis, but this could also be a by-product of selection on a different trait (16). Evidence from neurobiology suggests the sexes have both shared and dimorphic neural circuitry regulating sex-specific aggression (30, 31). Aggression may therefore be regulated by both sex-specific and shared genes.

Mating increases female aggression over food in *Drosophila melanogaster*, with mated females fighting for up to three times as long as unmated females of the same genotype (32). Like many post-mating responses (PMRs), the change in aggression after mating is stimulated by the transfer of sperm and at least one seminal fluid protein to females during mating (32). The strength of PMRs, including changes in remating and egg production rates, can vary because male genotype, condition, and age all influence the amount and content of ejaculate transferred to females (32, 33). Similarly, female genotype, condition, and age can also influence the strength and direction of PMRs because females differ in their response to male ejaculates. Thus the magnitude of increase in aggression after mating in females can evolve in response to the social environment (16).

We predicted that not only does genetic variation contribute to aggression in unmated females, but also to mating-induced aggression. Variation in this PMR may be due to differences in male ejaculates, female responses to ejaculates, or an interaction between the two. To measure genetic variation in aggression for males and females, and determine if aggression is correlated between the sexes, we measured the variation in male and female aggression among 10 isogenic strains of *Drosophila melanogaster* (Table 1). Nine lines were from the *Drosophila* Genome Resource Panel (DGRP). The DGRP is composed of lines derived from a single population that were inbred to near homozygosity (34). The lines have been used to determine the genetic basis of natural variation in male aggression (27). To measure lines which vary substantially in male and female aggression, we chose lines measured in Shorter et al. (27) with the highest and lowest rates of male aggression. We measured unmated male aggression, as well as unmated and mated female aggression in these lines. Using the same lines, we also determined if there is genetic variation in male induction of female aggression and female response to male ejaculates.

**Table 1.**
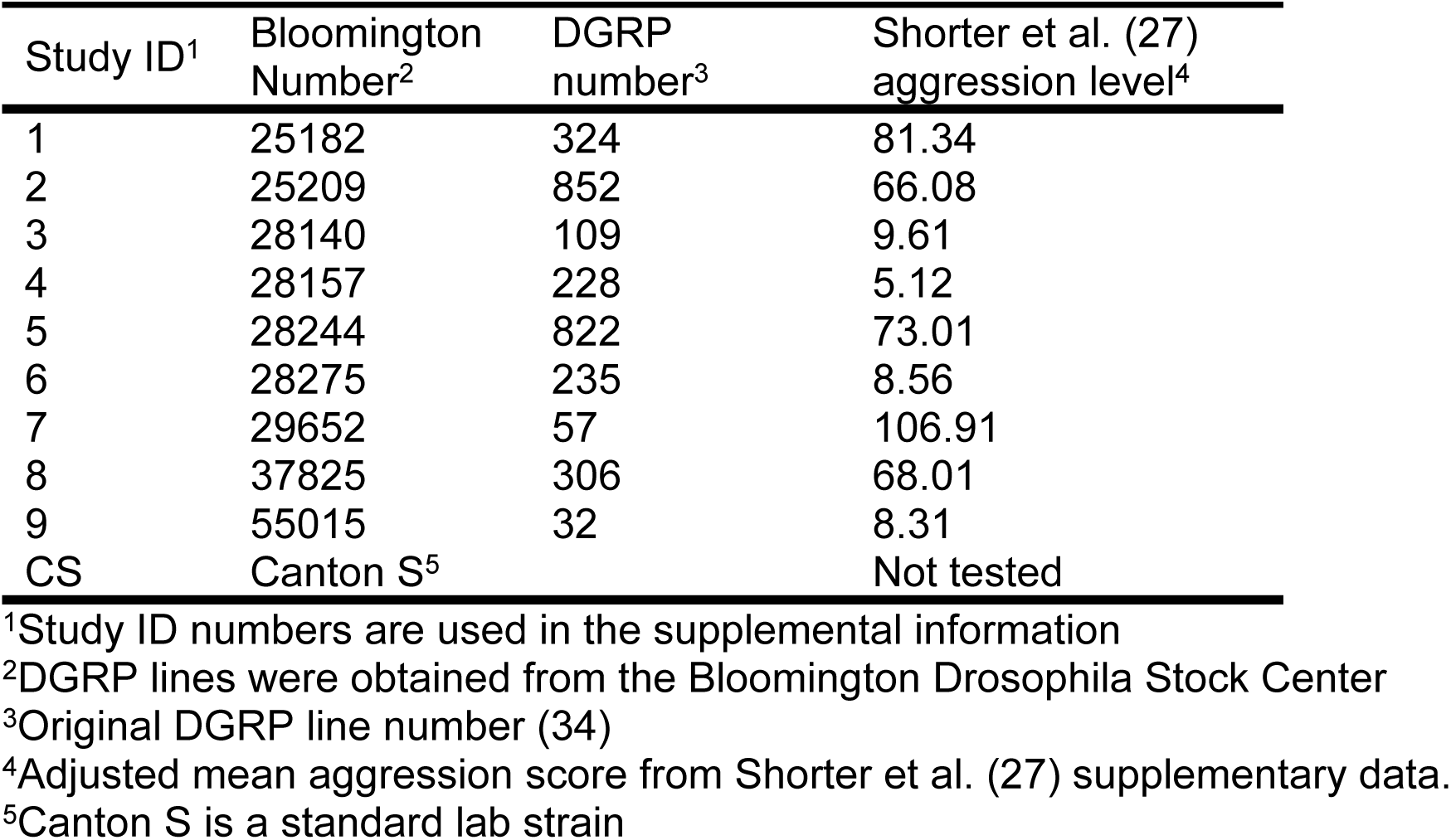
Strains used in this study.

| Study ID<sup>1</sup> | Bloomington Number<sup>2</sup> | DGRP number<sup>3</sup> | Shorter et al. (27) aggression level<sup>4</sup> |
| --- | --- | --- | --- |
| 1 | 25182 | 324 | 81.34 |
| 2 | 25209 | 852 | 66.08 |
| 3 | 28140 | 109 | 9.61 |
| 4 | 28157 | 228 | 5.12 |
| 5 | 28244 | 822 | 73.01 |
| 6 | 28275 | 235 | 8.56 |
| 7 | 29652 | 57 | 106.91 |
| 8 | 37825 | 306 | 68.01 |
| 9 | 55015 | 32 | 8.31 |
| CS | Canton S<sup>5</sup> |  | Not tested |

<sup>1</sup>Study ID numbers are used in the supplemental information
<sup>2</sup>DGRP lines were obtained from the Bloomington Drosophila Stock Center
<sup>3</sup>Original DGRP line number (34)
<sup>4</sup>Adjusted mean aggression score from Shorter et al. (27) supplementary data.
<sup>5</sup>Canton S is a standard lab strain

## Materials and Methods

### Fly culturing and strains

The stocks used in this study (Table 1) were obtained from the Bloomington Stock Center and are a subset of the Drosophila Genetics Resource Panel (DGRP, 34). We chose four of the lowest male aggression lines and the five highest male aggression lines as determined previously (27). The 10^th^ strain (Table 1) was a standard lab strain, Canton S (CS).

Flies were maintained at 25°C on a 12:12 light:dark cycle. Fly stocks were standardized by placing two females and two males in a large vial (24mm d x 94mm h) with cornmeal molasses fly food. Flies were sexed upon eclosion and marked with a dot of acrylic paint of one of two colors on the thorax to allow tracking of focal and competitor individuals in all assays. The standard competitor in all assays was from the CS strain. Color pairs varied among experiments but did not vary within experiments. Colors alternated between the focal individual and the CS competitor. Flies were kept individually in small vials (16.5mm d x 95mm h) with fly food until used in an assay. All flies were 3-6 days old when assayed.

### Aggression assays

Before an aggression assay, flies were starved individually in an empty small vial with a moist cotton plug (to avoid desiccation) for two hours to increase motivation to fight over food (28). A pair of flies, the focal fly and the standard competitor, were placed by aspiration in an aggression arena. An aggression arena was a well of a 12-well tissue culture dish (22 mm d x 20 mm h). In the arena was placed a food platform made from the lid of a 1.5 ml microcentrifuge tube filled with fly food and topped with a small amount of active yeast. Food platforms were only used once; tissue culture dishes were washed with mild soap and allowed to dry between assays. The well was covered with a transparent acetate sheet.

Assays were recorded using a Veho VMS-004 Discovery Deluxe USB Microscope camera connected to a PC. The recording was started before a focal individual and the standard competitor of the same sex were aspirated into the arena. The temperature and time of day were noted. Recordings continued for 30 minutes. Video recordings were saved for analysis once all assays in an experiment were completed.

### Male-male aggression

To determine if males vary in aggression, unmated males were assayed by placing one focal male (from the list in Table 1) with a CS male to control for variability in opponents. Ten assays were done for each line. Assays proceeded as above, starting with the starvation step.

### Unmated and mated female-female aggression

Unmated females were assayed in the same way as unmated males. Ten assays were done for each line. To generate mated females, one day before the assay the female of the focal line, and a CS female were individually placed with an unmated CS male and observed until mating occurred or two hours elapsed, whichever occurred first. Females that did not mate were discarded. The following day, the females were assayed as above, starting with starvation. Ten assays were performed for each line.

### Female aggression variation induced by males

To determine if males vary in their ability to manipulate female aggression, CS females were mated to males from each line and competed against CS females mated to CS males. Matings occurred as in the mated female assays above. Aggression assays proceeded as above, starting with starvation. Ten assays were done for each male line.

### Scoring and data analysis

All recordings were scored blind to treatment for the presence of aggressive behaviors using BORIS (35). All recordings were scored for the behaviors present as defined in Chen et al. (19) for males and Nilsen et al. (17) for females. Scored male behaviors included wing threats, lunging, chasing, boxing and tussling. Female behaviors including fencing and headbutts. For continuous behaviors, such as fencing, the duration of the behavior was recorded. The rate of aggressive behaviors was calculated and the number of events divided by the recording length.

All analyses were conducted in R v4.5.1 (36). We used generalized linear mixed effects models with negative binomial distributions to analyze the effects of DGRP line on focal and opponent individuals using the glmmTMB package (37). We checked model fit and assumptions using DHARMa (38). We established significance using the ‘Anov’ function from the car package (39). We calculated estimated marginal means and performed pairwise comparisons (with estimated marginal means adjusted for multiple comparisons with a Tukey adjustment) using the emmeans package (40) and plotted using the ggeffects (41) and patchwork packages (42). For female experiments, we used the number of headbutts performed by each individual fly as the response variable. For male analyses, we calculated the total number of aggressive events by adding together the number of boxing bouts, chases, headbutts, kicks, and wing threats. We then used the total number of aggressive events as the response variable in our male model. In all models, we included focal strain and temperature as fixed effects, with date and time since lights on as random effects. For the mated DGRP female experiment, we also included scorer as a fixed effect as there were multiple scorers. To investigate whether strains differed in their response to mating, we ran a model with data from both the unmated DGRP and mated DGRP female experiments. We used number of headbutts as the response variable. We included focal strain, mating status and their interaction as explanatory variables (to allow the change in aggression after mating to vary in magnitude between strains), as well as temperature as a fixed effect and date and time since lights on as random effects.

We investigated if Canton-S opponents changed their aggression based on the strain of the focal individual using generalized linear models with a negative binomial distribution. We used the number of headbutts by Canton-S individuals as the response variable in all female models, as well as including date and time since lights on as random effects. For DGRP females, we fit one model with both unmated and mated females included. As explanatory variables, we included the number of headbutts by the focal individual, mating status, the interaction between the two, focal strain and temperature as fixed effects. For the experiment with Canton-S females mated to DGRP males, our explanatory variables were the number of focal headbutts, strain of the male the focal female had mated with, and temperature as fixed effects. For DGRP males, we used the total number of aggressive events by the CS male as the response variable. We included the total number of aggressive events by the focal male, the strain of the focal male and temperature as fixed effects, with date and time since lights on as random effects.

To investigate whether there was a correlation between males and females within lines, we used model means for each line taken from our initial models (first paragraph above) for unmated DGRP females, mated DGRP females, and DGRP males. We then put in each model mean as the value for each strain and ran linear models to test for a correlation in aggression between the sexes for both unmated and mated females. To compare amount of variation between our different treatments (i.e. unmated DGRP females, mated DGRP females, mated Canton-S females, and males), we calculated the coefficient of variation (standard deviation/mean) for the focal flies for each experiment. We used the R package cvequality (43) to test for significant differences using the asymptotic test for equality of CVs in k populations.

Shorter et al. (27) measured aggression in males in 200 DGRP lines using a different assay. We asked if their aggression scores were correlated with our male aggression scores. To compare results from our male experiment with those of Shorter et al. (27). we took the model means from our model investigating the total number of aggressive events for males, which accounted for random effects of date and video recording time and used these model means as our response variable in a linear model. We included the total aggressive events for each line from Shorter et al. (27) as the explanatory variable.

## Results

### DGRP males

Focal strain had a significant effect on the total number of male aggressive events (Χ_29_ = 24.56, *P* = 0.004; Fig 1). Strain 306 was significantly different from strain 109 and the Canton-S strain but no other pairwise comparisons differed (S1 Table). Temperature did not affect the number of aggressive events (Χ_21_ = 0.02, *P* = 0.9).

**Fig 1.**
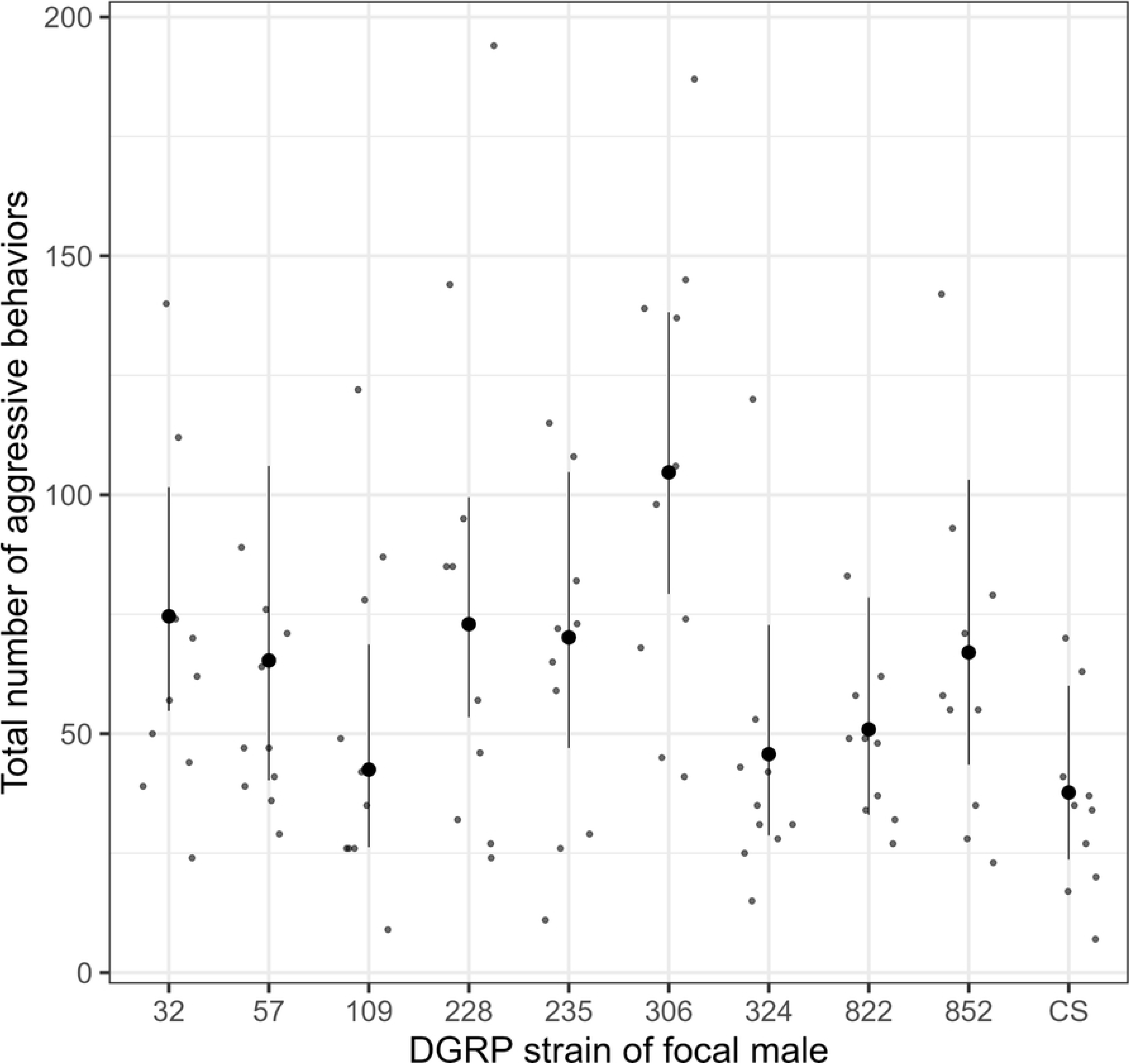
Male aggression differed among lines. Small points represent individual focal flies. Large points represent model means, with error bars representing 95% confidence intervals.

The number of aggressive events in our male experiment were not correlated with the aggression scores of Shorter et al. (27; Fig 2, F_1,7_ = 0.008, *P* = 0.93). When we focused on two separate high-intensity forms of aggression, boxing and chasing, we similarly found no correlations (Boxing: F_1,7_ = 0.11, *P*= 0.75; Chases: F_1,7_ = 0.07, *P* = 0.80).

**Fig 2:**
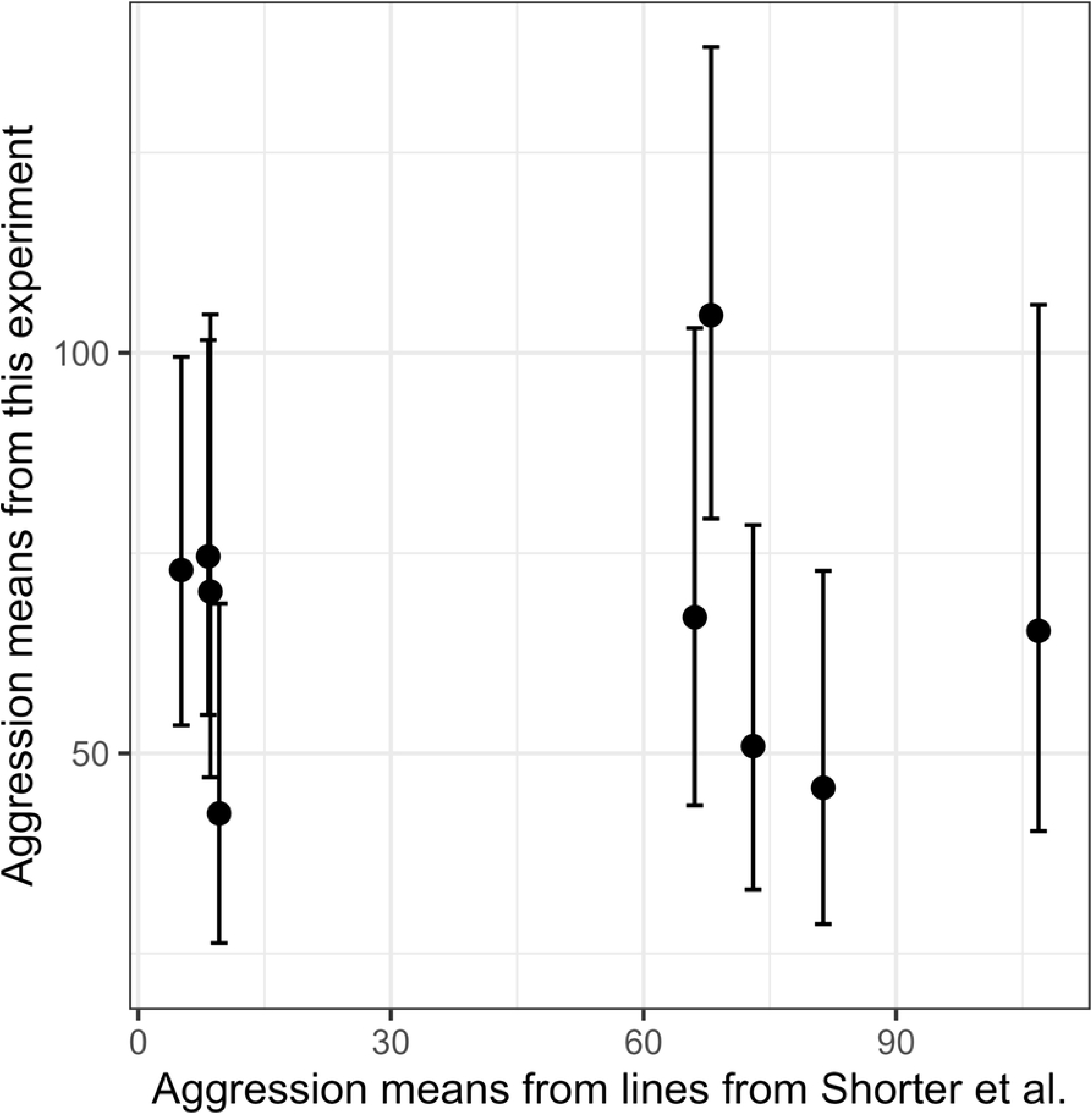
Male aggression measured in this study was not correlated with male aggression in a previous study (27). Points represent means for total events of male aggression taken from (27) and from the above male model for the current study. Vertical error bars represent the 95% confidence interval around the means from the current study.

### Females

The focal strain did not affect the number of headbutts performed by unmated females (Χ^2^_9_= 15.56, *P* = 0.08), nor did temperature have an effect (Χ^2^_1_ = 1.04, *P* = 0.31). Focal strain had a significant effect on the number of headbutts performed by mated females (Χ^2^_9_ = 29.36, *P* < 0.0001; Fig 3). Pairwise comparisons revealed that most strains were not different with the exception that strain 852 was significantly more aggressive than strains 32, 228, and 852 (S2 Table). This was the only experiment in which two people scored the videos; scorer did not have an effect (Χ^2^_1_ = 1.01, *P* = 0.32), nor did temperature (Χ^2^_1_ = 0.01, *P* = 0.91).

**Fig 3:**
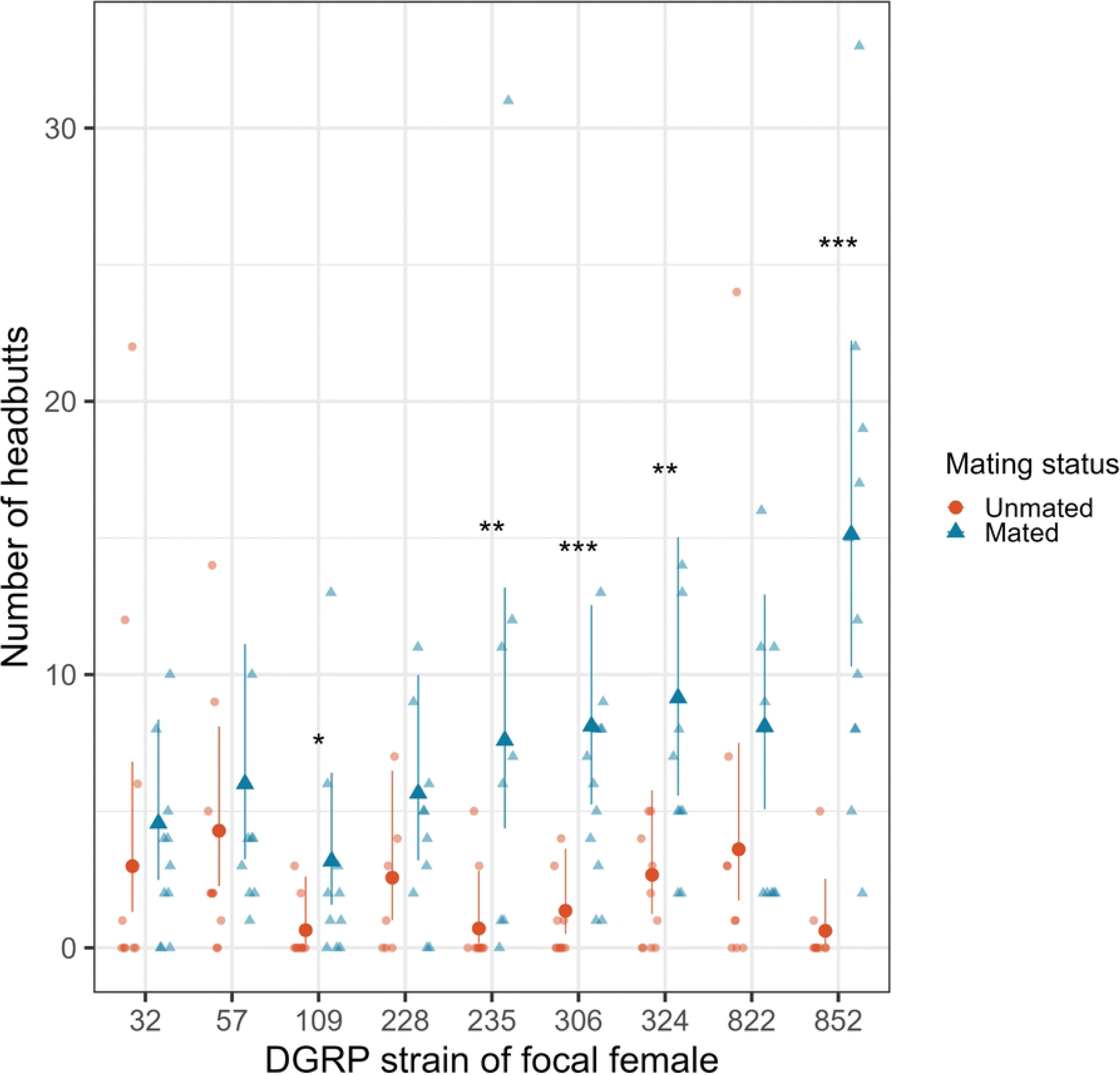
Lines differed in how much female aggression changed in response to mating. Small points represent individual focal flies. Large points represent model means, with error bars representing 95% confidence intervals. Asterisks indicate significant differences between unmated and mated females within a line (from pairwise comparisons with a Tukey multiple comparison adjustment). * = P <0.05, ** = P < 0.01, *** = P < 0.001.

### Effects of mating for DGRP females

Focal strain and mating status had a significant interaction influencing the number of headbutts performed by females (Fig 3, Χ^2^_8_ = 18.16, *P* = 0.02; S3 Table). Temperature did not have an effect (Χ^2^_1_ = 0.07, *P* = 0.80). Five lines increased aggression after mating, but the magnitude of that change was variable, ranging from three times as many headbutts (Line 324: Unmated = 2.25 ± 0.96, Mated = 7.47 ± 0.99) to 24 times as many (Line 852: Unmated = 0.53 ± 0.39, Mated = 12.29 ± 2.6). The remaining lines did not increase, nor decrease, aggression after mating. Unmated and mated headbutts were not correlated across lines (F_1,7_ = 0.58, *P* = 0.47; S1 Fig).

### CS females mated to DGRP males

To determine if male strain affected female aggression, CS females were mated to males of each DGRP strain, as well as CS. Focal male strain did not affect the level of aggression in Canton-S females (Fig 4, Χ^2^_9_ = 8.36, *P* = 0.50), nor did temperature (Χ^2^_1_ = 1.40, *P* = 0.24).

**Fig 4.**
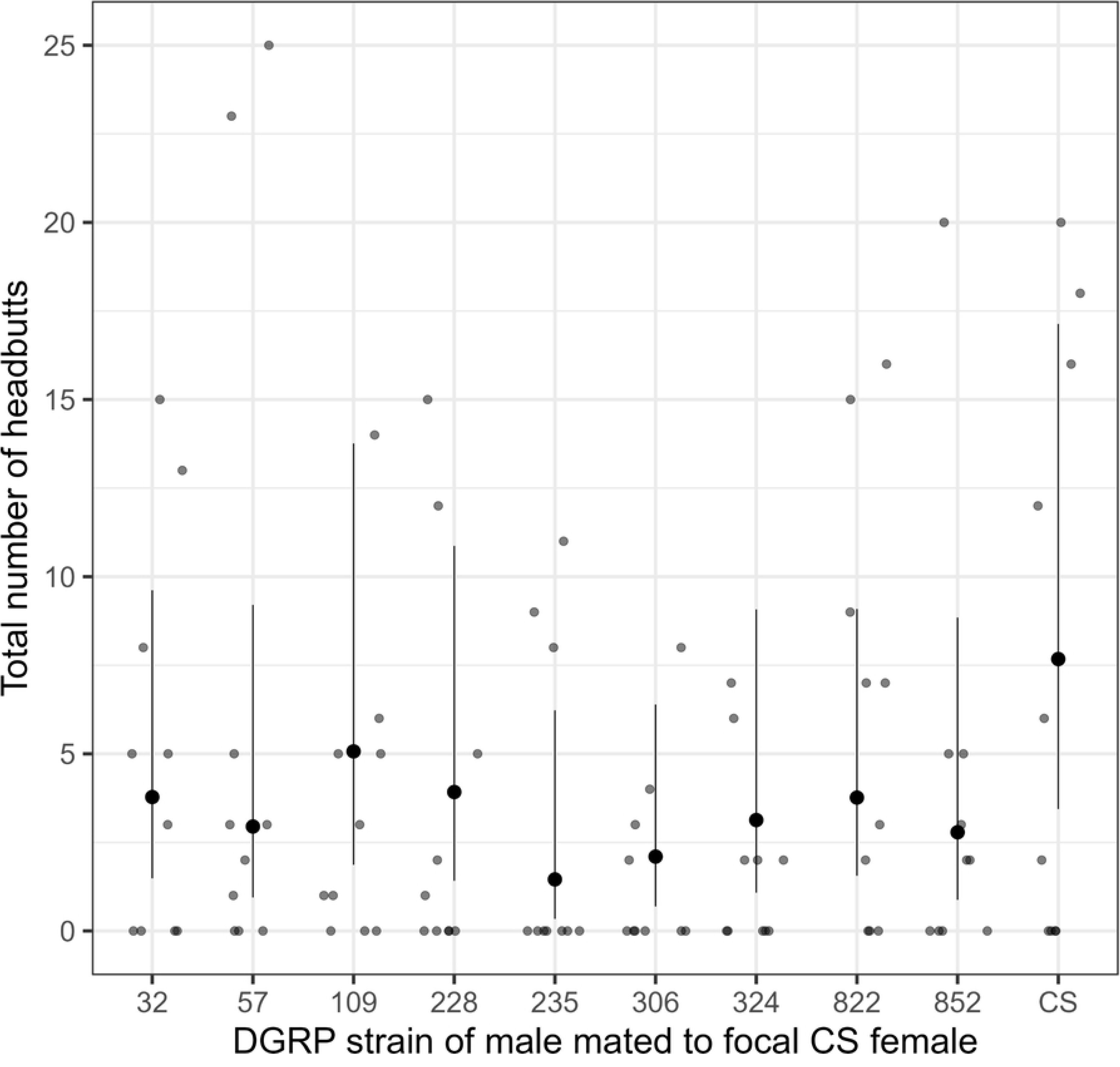
Mated aggression did not change with male strain. CS females were mated to each line and aggression was measured. Small points represent individual focal flies. Large points represent model means, with error bars representing 95% confidence intervals

### Correlations between CS opponents and focal individuals

The total number of aggressive events by the focal male and the number from their CS opponent were correlated (Fig 5A, Χ^2^_1_ = 37.43, *P* < 0.001). Neither focal strain (Χ^2^_9_ = 6.65, *P* = 0.67) nor temperature (Χ^2^_1_ = 2.08, *P* = 0.15) affected total male aggressive events. Similarly, the number of boxing bouts started by the focal male was positively correlated with the number started by their CS opponent (Χ^2^_1_ = 8.54, *P* = 0.003) with no effect of focal strain (Χ^2^_9_ = 4.05, *P* = 0.91), or of temperature (Χ^2^_1_ = 0.35, *P* = 0.55). In contrast, the number of focal chases and the opponent CS opponent chases were not correlated (Χ^2^_1_ = 2.87, *P* = 0.09); nor were there any effects of focal strain (Χ^2^_9_ = 15.34, *P* = 0.08) or temperature (Χ^2^_1_ = 0.001, *P* = 0.97).

**Fig 5:**
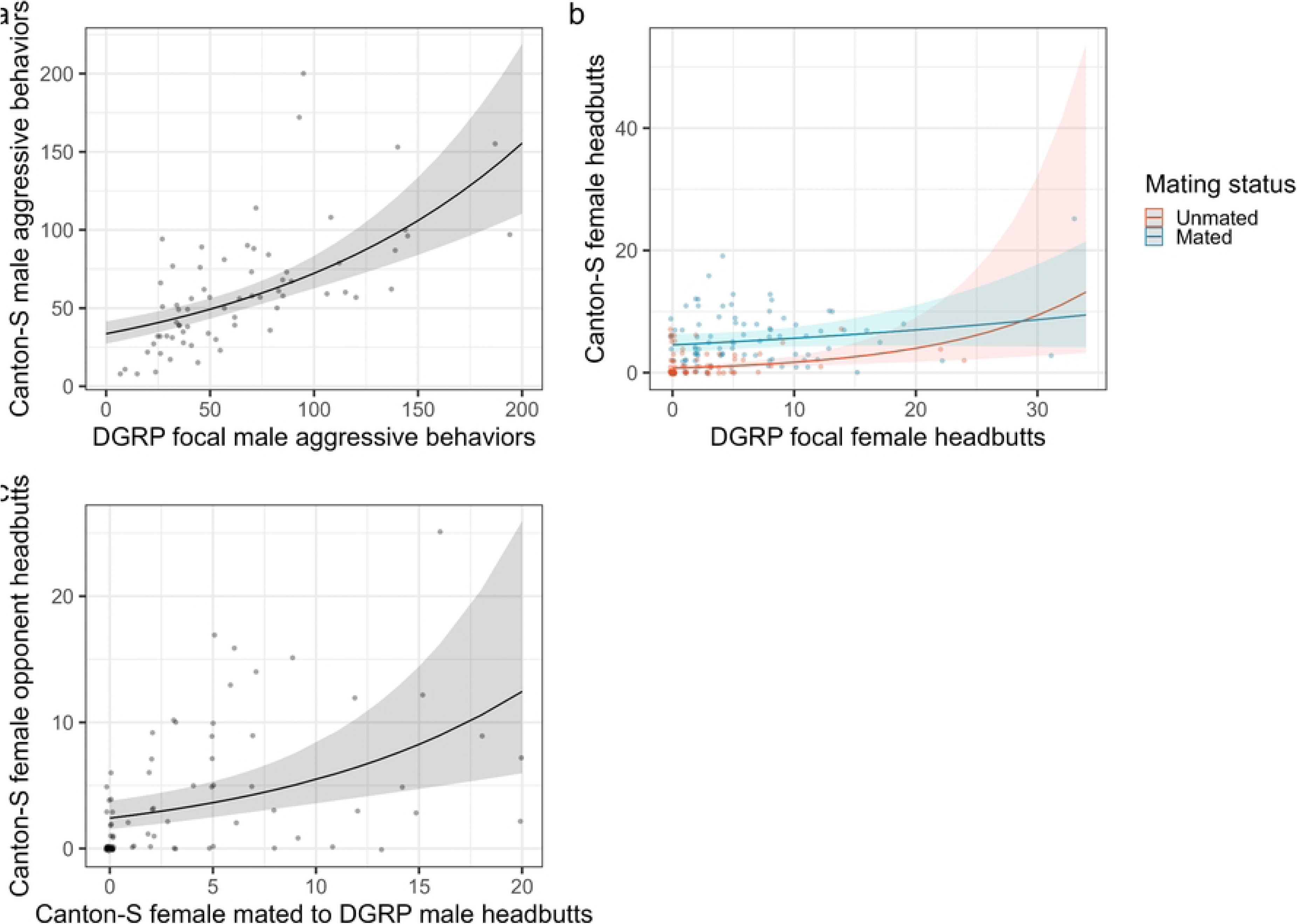
The number of aggressive behaviors performed by the focal individual was correlated with the behavior of the opponent (control) individual in most experiments. Correlations between the number of aggressive behaviors by the focal individual and their opponent in A) DGRP males; B) DGRP females (unmated in red, mated in blue); and C) CS females mated to DGRP males. Small points represent individual dyads. The lines indicate model predictions of the correlation with the shaded error indicating a 95% confidence interval.

In the experiments with DGRP females, mating status and focal headbutts had a significant interaction with the number of headbutts performed by CS opponents (Fig 5B, Χ^2^_1_ = 5.26, *P* = 0.02). A significant positive relationship existed between focal and CS opponent headbutts in unmated females (Estimated marginal trend = 0.085 ± 0.023), but not in mated females (0.022 ± 0.015).

A significant positive relationship existed between focal headbutts and opponent headbutts in Canton-S females mated to DGRP males (Fig 5C, Χ^2^_1_ = 13.49, *P* < 0.001). Neither strain of the male (Χ^2^_9_ = 7.7, *P* = 0.56), nor temperature (Χ^2^_1_ = 0.31, *P* = 0.58) affected the relationship.

### Correlations between males and females

To determine if males and females have a common basis for aggression, we compared total male aggression and female headbutts. We found no correlations across DGRP lines between the number of headbutts by unmated females and the number of aggressive encounters by males (Fig 6A, F_1,7_ = 0.12, *P* = 0.74). The number of headbutts by mated females and the number of aggressive encounters by males were similarly not correlated (Fig 6B, F_1,7_ = 0.08, *P* = 0.79).

**Fig 6.**
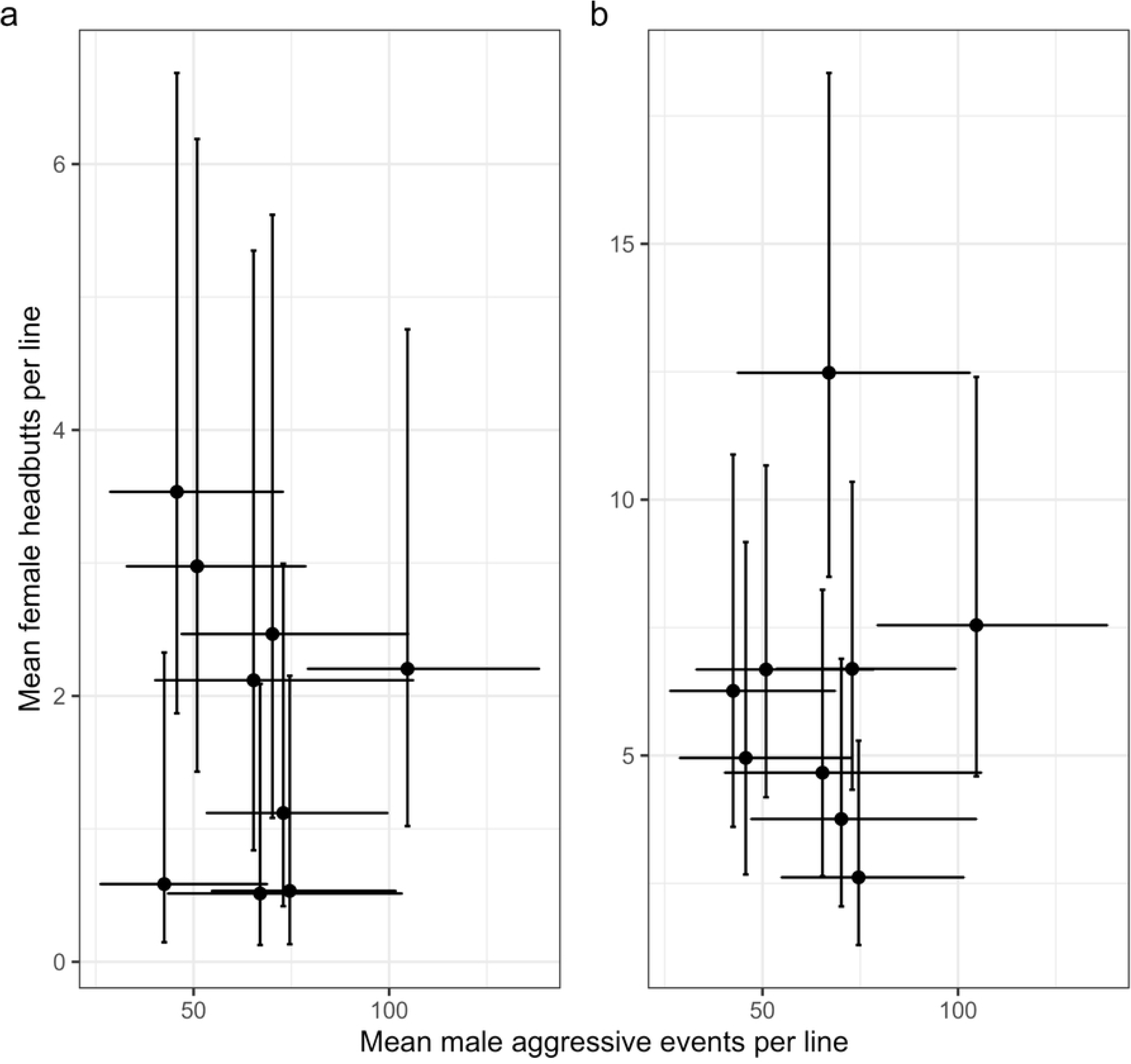
Male and female aggression were not correlated across the DGRP lines. Point represent means for female and male aggression taken from the model analyzing DGRP female aggression. Error bars represent the 95% confidence interval. A). Unmated females. B). Mated females

To compare variation between our different groups, we calculated the coefficients of variation for each different set of focal flies. We found significant differences between males and all groups of females (D’AD = 102.27, *P* < 0.0001, CV_males_ = 0.63). Unmated DGRP females had more variation than mated DGRP females (D’AD = 7.17, *P* = 0.007, CV_unmated_ = 1.98, CV_mated_ = 0.97). There was no significant difference between unmated DGRP females and mated Canton-S females (D’AD = 1.51, *P* = 0.22, CV_Canton-S_ = 4.15) or between mated DGRP females and mated Canton-S females (D’AD = 2.83, *P* = 0.09).

## Discussion

### Males

As expected, males varied in their levels of aggression; we saw a similar magnitude of variation as previous studies (Coefficient of variation: our study = 0.31, Shorter et al. (27) = 0.55, D’AD = 2.09, *P* = 0.15). However, our measures of aggression did not correlate with measures from the same lines in Shorter et al. (27). Our method of scoring aggression, although similar to most male aggression studies (44, 45), was substantially different from that of Shorter et al. (27). First, we housed males individually after eclosion whereas Shorter et al. (27) maintained males in mixed sex groups. Social experience affects behaviors (reviewed in 46) including aggression (47–49), thus previous encounters are likely to shape the results of Shorter et al. (27). Notably, aggression increases with isolation in *D. melanogaster* males (50) and females (23). Differences between Shorter et al. (27) and our results may reflect genetic variation in response to social isolation among the lines as well as genetic variation in aggression.

Another major difference in the way we scored aggression, versus the method of Shorter et al. (27) is that we scored focal individuals with a single standardized opponent (a CS male) whereas Shorter et al. (27) scored the number of aggressive interactions in groups of eight males of the same strain. As we observed, males of our standard CS line modified their behavior in response to their opponent with increased aggression towards a DGRP male relative to a CS male. Aggression is not only influenced by the individual, but also by the social environment thus indirect genetic effects (IGEs) are likely at play because an individual’s behavior can affect and be affected by the behaviors of the interacting individual (51, 52). Thus, the genotype of the competitor affected the behavior of the CS male. If we competed DGRP males against males of their own genotype, we might have seen greater variation in aggression among DGRP with males fighting with males of similar aggression levels, either amplifying or suppressing aggression levels relative to CS. The addition of extra males (27) also influences rates of aggression and the genotype of those additional males affects rates of aggression in the population (51). Shorter et al. (27) may therefore have greater effects of strain compared to us due to many males of the same genotype interacting.

### Females

Genetic variation in female aggression is not well-studied. We found that the DGRP lines varied in aggression in mated females. The coefficients of variation for females were more variable in all groups of females compared to males. Unmated DGRP females had the highest CV (1.98), followed by CS females mated to DGRP males (1.42), DGRP females mated to CS males (0.97), and finally DGRP males (0.63). This suggests high levels of variation in female aggression, even if this is not among genetic lines (as we found no differences between lines in unmated DGRP females or mated CS females). These results suggest that females have more standing genetic variation in aggression in females than males, which may reflect weak selection on female aggression leading to genetic drift and slow purging of genetic variants related to aggression (53). Females are unlikely to remain unmated for long in wild populations, leading to few opportunities for selection on unmated female aggression. Unmated females may not need to compete for access to resources, leading to weak selection for aggression, which would explain the low levels of aggression in unmated females and the relatively large amount of variation.

The post-mating change in aggression varied by line with five lines increasing aggression and four lines remaining unchanged after mating. The increase in aggression associated with mating has been observed in other lines of *Drosophila melanogaster* (20, 32) but not in other species, which tend to decrease aggression post-mating, if they change at all (54). Our results indicate that mating-induced change in aggression and its magnitude varies within a species.

During copulation, males transfer ejaculates containing sperm and seminal fluid proteins that interact with receptors and proteins in the female reproductive tract to alter female physiology and behavior (e.g. 55-57). Both variation among males in the composition of their ejaculates and variation in female reproductive tract traits can influence the strength of female post-mating responses (PMRs, 58). Male and female reproductive traits may coevolve, leading to interactions between male and female genotypes that shape female PMRs (59). For example, females from experimentally evolved lines mated with standard males do not differ in mating-induced aggression among lines, but when mated with males from their own lines, lines differed in response to mating (16).

Because of the interactions between males and females during mating, differentiating between male-induced effects and female responses to male ejaculates is difficult. With our experimental design, we could distinguish between male effects on females and female responses to mating. By mating standard CS females to DGRP males, we found no evidence that males differ in their ability to stimulate aggression in females. In another study, standard females had the same response (in egg-laying and remating rate) to mating independent of the line of the male to whom she was mated; however, females varied among lines in their response to mating and sex-peptide injection, implying that females vary in receptivity to male ejaculates (59). We observed the same phenomenon: variation in DGRP line for female response to mating, but lack of variation within a line.

Previous work suggested that the strength of mating-induced aggression was related to the amount and/or quality of sperm females receive during mating (60). Because all CS males were the same age and unmated before the experiment, all mated females should have received similar amounts of sperm and seminal fluid proteins and enough of both to induce the maximum post-mating increase in aggression. Variation in aggression post-mating may exist for two reasons: 1) lines vary in the maximum level of aggression they can reach regardless of how much ejaculate they receive. Because females are more aggressive after mating, they only hit the aggression ceiling after mating, so we only observe differences between strains in the mated females and how much they change relative to unmated females. 2) the lines may vary in sperm ejection or sperm storage traits (61). Because the quantity and quality of sperm influence rates of aggression, if females differ in how much sperm they store after mating, this may influence their subsequent rate of mating-induced aggression.

### Relationship between male and female aggression

Aggression of males and females in the DGRP lines was not correlated, implying a different genetic basis to the behavior for the sexes. The lack of correlation might be predicted from several lines of evidence. First, the outcomes of fighting are different between males and females: male fights can result in the formation of dominance hierarchies (17) but this is not an outcome of female fights. Males escalate aggression over fights whereas females engage in low intensity headbutts (17, 23).

Second, the neurological basis of aggression is both similar and different between males and females. Males and females share a common upstream neuron that induces aggressive behavior through sexually dimorphic downstream neurons (30, 31). Females retain the structures necessary for male aggression because genetic masculinization results in male-typical aggressive behaviors when facing a male opponent, although they exhibit female-typical aggressive behaviors against a female opponent (62). Thus, males and females have both common and differentiated neurological underpinnings to aggression.

Finally, selection experiments on male aggression behaviors are mixed with studies finding either increased female aggression when male aggressive behavior is selected (28) or an absence of a correlated response (29). Although male aggression has been genetically manipulated and gene expression associated with aggression has been measured (e.g. 7), similar studies have not been performed with female aggression. Gene expression studies associated with female behavior are needed. In addition, experiments selecting for female aggression should be done to determine if a correlated response occurs in males.

### Response to opponent

In all experiments, except those with DGRP mated females, the number of aggressive behaviors by the Canton-S opponent was positively correlated with that of the focal fly. These results again suggest indirect genetic effects are influencing rates of aggression as the genotype of the focal fly shapes the aggression rate of the competitor fly (51, 63). These relationships may be the result of two non-mutually exclusive processes. First, initiating an attack and responding to being attacked may represent two different components of aggression (15, 64). If a CS fly is facing a more aggressive opponent, it will be attacked more often and therefore have more opportunities to respond to attacks than if it was facing a less aggressive opponent. The two behaviors, initiating attached and responding to attacks, are likely genetically correlated, but respond differently to selection (15). The trends in Fig 5 tell us that overall focal flies attacked more often than CS flies (the slope of the line does not follow a 1:1 ratio), which suggests that CS flies are only responding to some attacks by focal flies. However, when we look at the raw data (Fig. 5), the focal fly is not always the most aggressive individual in a dyad, which does not fit with CS flies responding to more aggressive focal fly attacks. We do not have information on which fly initiated encounters or the order of events, so we cannot determine if CS aggression occurs predominantly after the fly has been attacked.

A second explanation for the positive correlation is that *D. melanogaster* flies are possibly using a form of mutual assessment in their aggressive encounters. Theoretical and empirical work has demonstrated that the duration and maximum intensity of a fight is determined by the eventual loser – the decision to retreat ends the fight (65). Individuals may decide when to end a fight based on an assessment of their own resource holding potential (self-assessment) or also incorporate information about their opponent’s resource holding potential (RHP; mutual assessment). Although we do not have data on the RHP of flies in our experiments, previous work on females suggests that flies that initiate attacks win more encounters (21), suggesting that rate of aggression may be a useful indicator of the eventual winner in a contest. Our results are consistent with previous work on males, which suggests *D. melanogaster* may use some form of mutual assessment (19). We used CS flies raised at standard density as the opponents for all experiments, which should have limited variation in RHP among CS opponents, but not eliminate it entirely. However, we do not have information on the RHP of flies, the winners of these encounters, or patterns of escalation which would help us to establish which contest assessment strategies are most likely in this species.

## Conclusion

As expected, both males and females had genetic variation for aggression. Additionally, females varied in their response to mating, though males did not vary in their ability to induce a post-mating response. Aggression was not correlated between males and females, suggesting sex-specific genetic architecture. Because most studies of genes affecting aggression have focused on males, studies of the genetics of female aggression are needed. Studies should sample more lines than we studied here, use RNA-seq and GWAS to identify candidate genes, and perform gene manipulation experiments targeted at females. Such studies will improve our understanding of aggression by identifying loci underlying female-specific aggression as well as the plastic effects of mating on aggression. Studying natural variation in behaviors within and across populations will give us important insights into both the mechanistic basis and evolutionary function of aggression.

## Acknowledgments

We thank Carine Tabak for the producing preliminary data that inspired the full study. Cole VandeVelde, Sarah Strong, Madeline Halabi, and Oluwafifeyemi Babtope-Ojo all contributed to data generation.

## Supporting Information

**S1 Table. Pairwise comparisons for DGRP male aggressive behaviors**

**S2 Table. Pairwise comparisons for mated DGRP females**

**S3 Table. Means for unmated and mated DGRP females, with effects of mating**

**S1 Fig. Absence of correlation between unmated and mated female headbutts across lines.** Points represent means for female and male aggression taken from the model analyzing DGRP female aggression. Error bars represent the 95% confidence interval.

## References

1. Clutton-Brock TH, Huchard E. Social competition and selection in males and females. Phil Trans R Soc B. 2013;368(1631):20130074. doi: 10.1098/rstb.2013.0074

2. Georgiev AV, Klimczuk ACE, Traficonte DM, Maestripieri D. When violence pays: A cost-benefit analysis of aggressive behavior in animals and humans. Evolutionary Psychology. 2013;11(3):678–99. doi: 10.1177/147470491301100313

3. Sih A, Bell A, Johnson JC. Behavioral syndromes: an ecological and evolutionary overview. Trends Ecol Evol. 2004;19(7):372–8. doi:10.1016/j.tree.2004.04.00

4. Anholt RRH, Mackay TFC. Genetics of aggression. Annual Review of Genetics. 2012;46:145–64. doi: 10.1146/annurev-genet-110711-155514

5. Hecht EE, Kukekova AV, Gutman DA, Acland GM, Preuss TM, Trut LN. Neuromorphological changes following selection for tameness and aggression in the Russian farm-fox experiment. Journal of Neuroscience. 2021;41(28):6144–56. doi: 10.1523/JNEUROSCI.3114-20.2021

6. Suzuki K, Okanoya K. Domestication effects on aggressiveness: Comparison of biting motivation and bite force between wild and domesticated finches. Behavioural Processes. 2021;193:104503. doi: 10.1016/j.beproc.2021.104503

7. Dierick HA, Greenspan RJ. Molecular analysis of flies selected for aggressive behavior. Nature Genetics. 2006;38(9):1023–31.

8. Hecht S, Wald G. The visual acuity and intensity discrimination of Drosophila. J Gen Physiol. 1934;17(4):517–47.

9. Archard GA, Braithwaite VA. Variation in aggressive behaviour in the poeciliid fish *Brachyrhaphis episcopi*: Population and sex differences. Behavioral Processes. 2011;86:52–7. doi: 10.1016/j.beproc.2010.09.002

10. Magurran AE, Seghers BH. Variation in schooling and aggression amonst guppy (*Poecilia reticulataI*) populations in Trinidad. Behaviour. 1991;118(3/4):214–34.

11. Yokel DA. Intrasexual aggression and the mating behavior of brown-headed cowbirds: their relation to population densities and sex ratios. Condor. 1989;91:43–51.

12. Kilgour RJ, McAdam AG, Betini GS, Norris DR. Experimental evidence that density mediates negative frequency-dependent selection on aggression. J Anim Ecol. 2018;87:1091–101. doi: 10.1111/1365-2656.12813

13. Clutton-Brock T, Huchard E. Social competition and its consequences in female mammals. J Zool. 2013;289(3):151–71. doi: 10.1111/jzo.12023

14. Ah-King M. The history of sexual selection research provides insights as to why females are still understudied. Nat Commun. 2022;13:6976. doi: 10.1038/s41467-022-34770-z

15. Hemmat M, Eggleston P. Competitive interactions in *Drosophila melanogaster*: recurrent selection for aggression and response. Heredity. 1988;60:129–37.

16. Bath E, Edmunds D, Norman J, Atkins C, Harper L, Rostant WG, et al. Sex ratio and the evolution of aggression in fruit flies. Proc R Soc London, Ser B. 2021;288(1947):20203053. doi: 10.1098/rspb.2020.3053

17. Nilsen SP, Chan Y-B, Huber R, Kravitz EA. Gender-selective patterns of aggressive behavior in *Drosophila melanogater*. Proc Natl Acad Sci USA. 2004;101(33):12343–7. doi: 10.1073/pnas.0404693101

18. Baxter CM, Barnett R, Dukas R. Aggression, mate guarding and fitness in male fruit flies. Anim Behav. 2015;109: 235–41. doi: 10.1016/j.an.behav.2015.08.023

19. Chen S, Lee AY, Bowens NM, Kravitz EA. Fighting fruit flies: a model system for the study of aggression. Proc Natl Acad Sci USA. 2002;99(8):5664–8. doi: 10.1073/pnas.082102599

20. Bath E, Biscocho ER, Easton-Calabria A, Wigby S. Temporal and genetic variation in female aggression after mating. PLoS ONE. 2020;15(4):e0229633. doi: 101371/journal.pone.0229633

21. Bath E, Morimoto J, Wigby S. The developmental environment modulates mating-induced aggression and fighting success in adult female *Drosophila*. Funct Ecol. 2018;32(11):2542–52. doi: 10.1111/1365-2435.13214

22. Billeter J-C, Jagadeesh S, Stepek N, Azanchi R, Levine JD. *Drosophila melanogaster* females change mating behaviour and offspring production based on social context. Proc R Soc London, Ser B. 2012;279(1737):2417–25. doi: 10.1098/rspb.2011.2676

23. Ueda A, Kidokoro Y. Aggressive behaviours of female Drosophila melanogaster are influenced by their social experience and food resources. Physiol Entomol. 2002;27:21–8.

24. White MA, Chen DS, Wolfner MF. She’s got nerve: roles of octopamine in insect female reproduction. J Neurogenet. 2021;35(3):132–53. doi: 10.1080/01677063.2020.1868457

25. Gaspar M, Dias S, Vasconcelos ML. Mating pair drives aggressive behavior in female *Drosophila*. Curr Biol. 2022;32(21):4734–42. doi: 10.1016/j.cub.2022.09.009

26. Hoffmann AA. Heritable variation for territorial succes in two *Drosophila melanogaster* populations. Anim Behav. 1988;36:1180–9.

27. Shorter J, Couch C, Huang W, Carbone MA, Peiffer J, Anholt RRH, et al. Genetic architecture of natural variation in *Drosophila melanogaster* aggressive behavior. Proc Natl Acad Sci USA. 2015;112:E3555–E63. doi: 10.1073/pnas.1510104112

28. Edwards AC, Rollman SM, Morgan TJ, Mackay TFC. Quantitative genomics of aggressive behavior in *Drosophila melanogaster*. PLoS Genet. 2006;2(9):e154. doi: 10.1371/journal.pgen.0020154

29. Penn JKM, Zito MF, Kravitz EA. A single social defeat reduces aggression in a highly aggressive strain of *Drosophila*. Proc Natl Acad Sci USA. 2010;107(28):12682–6. doi: 10.1073/pnas.1007016107

30. Chiu H, Hoopfer ED, Coughlan ML, Anderson DJ. A circuit logic for sexually shared and dimorphic aggressive behaviors in *Drosophila*. Cell. 2021;184(2):507–20. doi: 10.1016/j.cell.2020.11.048

31. Schretter CE, Aso Y, Robie AA, Dreher M, Dolan M-J, Chen N, et al. Cell types and neuronal circuitry underlying female aggression in *Drosophila*. eLife. 2020;9(1):e58942. doi: 10.7554/eLife.58942

32. Bath E, Bowden S, Peters C, Reddy A, Tobias JA, Easton-Calabra E, et al. Sperm and sex peptide stimulate aggression in female *Drosophila*. Nat Ecol Evol. 2017;1(6):0154. doi: 10.1038/s41559-017-0154

33. Avila FW, Sirot LK, Laflamme BA, Rubinstein CD, Wolfner MF. Insect seminal fluid proteins: identification and function. Ann Rev Ent. 2011;56:21–40. doi: 10.1146/annurev-ento-120709-144823

34. Mackay TFC, Richards S, Stone EA, Barbadilla A, Ayroles JF, Zhu D, et al. The *Drosophila melanogaster* genetic reference panel. Nature. 2012;482:173–8. doi: doi:10.1038/nature10811

35. Friard O, Gamba M. BORIS: a free, versatile open-source event-logging software for video/audio coding and live observations. Methods in Ecology and Evolution. 2016;7:1325–30. doi: 10.1111/2041-210X.12584

36. R Core Team. R: A language and environment for statistical computing. Viena, Austria: R Foundation for Statistical Computing; 2017.

37. McGillycuddy M, Popovic G, Bolker BM, Warton DI. Parsimoniously fitting large multivariate random effects in glmmTMB. J Stat Softw. 2025;112(1):1–19. doi: 10.18637/jss.v112.i01

38. Hartig F. DHARMa: Residual Diagnostics for Hierarchical (Multi-Level / Mixed) Regression Models. 2024.

39. Fox JH, Weisberg S. An R Companion to Applied Regression. 3rd ed. Thousand Oaks, CA: Sage Publications; 2018.

40. Lenth R, Piaskowski J. emmeans: Estimated Marginal Means, aka Least-Squares Means. 2.0.3 ed 2026.

41. Lüdecke D. ggeffects: tidy data frames of marginal effects from regeression models. Journal of Open Source Software. 2018;3(26):772. doi: 10.21105/joss.00772

42. Pedersen T. patchwork: The Composer of Plots. 1.3.2 ed 2025.

43. Marwick B, Krishnamoorthy K. cvequality: Tests for the Equality of Coefficients of Variation from Multiple Groups. R software package version 0.1.3 ed: Github; 2019.

44. Dankert H, Wang L, Hoopfer ED, Anderson DJ, Perona P. Automated monitoring and analysis of social behavior in *Drosophila*. Nat Methods. 2009;6(4):297–303. doi:10.1038/nmeth.1310

45. Leng X, Wohl M, Ishil K, Nayak P, Asahina K. Quantifying influence of human choice on the aautomated detection of *Drosophila* behavior by a supervised machine learning algorithm. PLoS ONE. 2020;15(12):e0241696. doi: 10.1371/journal.pone.0241696

46. Sokolowski MB. Social interaction in “simple” model systems. Neuron. 2010;65:780–94. doi: 10.1016/j.neuron.2010.03.007

47. Nandy B, Dasgupta P, Halder S, Verma T. Plasticity in aggression and the correlated changes in the cost of reproduction in male *Drosophila melanogaster*. Anim Behav. 2016;114:3–9. doi: 10.1016/j.anbehav.2016.01.019

48. Chen M, Sokolowski MB. How social experience and environment impacts behavioural plasticity in *Drosophila*. Fly. 2022;16(1):68–84. doi: 10.1080/19336934.2021.1989248

49. Yurkovic A, Wang O, Basu AC, Kravitz EA. Learning and memory associated with aggression in *Drosophila melanogaster*. Proc Natl Acad Sci USA. 2006;103(46):17519–24. doi: 10.1073 pnas.0608211103

50. Wang L, Dankert H, Perona P, Anderson DJ. A common genetic target for environmental and heritable influences on aggressiveness in *Drosophila*. Proc Natl Acad Sci USA. 2008;105(15):5657–63. doi: 10.1073/pnas.0801327105

51. Saltz JB. Genetic composition of social groups influences male aggressive behaviour and fitness in natural genotypes of *Drosophila melanogaster*. Proc Natl Acad Sci USA. 2013;280:20131926. doi: 10.1098/rspb.2013.1926

52. Wilson AJ, Gelin U, Perron M-C, Réale D. Indirect genetic effects and the evolution of aggression in a vertebrate system. Proc R Soc London, Ser B. 2009;276:533–41. doi: 1098/rspb.2008.1193

53. Johnson OL, Tobler R, Schmidt JM, Huber CD. Fluctuating selection and the determinants of genetic variation. Trends in Genetics. 2023;39(6):491–504. doi: 101016/j.tig.2023.02.004

54. Bath E, Gleason JM. Is variation in female aggressiveness across Drosophila species associated with reproductive potential? Proc R Soc London, Ser B. 2025;292:20242301. doi: 10.1101/2024.08.27.609931

55. Chapman T, Herndon LA, Heifetz Y, Partridge L, Wolfner MF. The Acp26Aa seminal fluid protein is a modulator of early egg hatchability in *Drosophila melanogaster*. Proc R Soc London, Ser B. 2001;268:1647–54. doi: 10.1098/rspb.2001.1684

56. McGraw LA, Gibson G, Clark AG, Wolfner MF. Genes regulated by mating, sperm, or seminal proteins in mated female *Drosophila melanogaster*. Curr Biol. 2004;14:1509–14. doi: 10.1016/j.cub.2004.08.028

57. Wolfner MF. Battle and ballet: molecular interactions between the sexes in Drosophila. Journal of Heredity. 2009;100(4):399–410. doi: doi:10.1093/jhered/esp013

58. McDonough-Goldstein C, Pitnick S, Dorus S. *Drosophila* female reproductive glands contribute to mating plug composition and the timing of sperm ejection. Proc R Soc London, Ser B. 2022;289(1968):20212213. doi: 10.1098/rspb.2021.2213

59. Wensing K, Fricke C. Divergence in sex peptide-mediated female post-mating responses in *Drosophila melanogaster*. Proc R Soc London, Ser B. 2018;285(1886):20181563. doi: 10.198/rspb.2018.1563

60. Bath E, Buzzoni D, Ralph T, Wigby S, Sepil I. Male condition influences female post mating aggression and feeding in *Drosophila*. Funct Ecol. 2021;35:1288–98. doi: 10.1111/1365-2435-13791

61. Lüpold S, Reil JB, Manier MK, Zeender V, Belote JM, Pitnick S. How female × male and male × male interactions influence competitive fertilization in *Drosophila melanogaster*. Evolution Letters. 2020;4-5:416–29. doi: 10.1002/evl3.193

62. Monyak RE, Golbari NM, Chan Y-B, Pranevicius A, Tang G, Fernádez MP, et al. Masculinized *Drosophila* females adapt their fighting strategies to their opponent. J Exp Biol. 2021;224:jeb238006. doi: 10.1242/jeb.238006

63. Girardeau AR, Enochs GE, Saltz JB. Evolutionary feedbacks for *Drosophila* aggression revealed through experimental evolution. Proc Natl Acad Sci USA. 2025;122(17):e2419068122. doi: 10.1073/pnas.2419068122

64. Mather K, Caligari PDS. Pressure and response in competitive interactions. Heredity. 1983;51(2):435–54.

65. Maynard Smith J, Price GR. The logic of animal conflict. Nature. 1973;246:15–8.

